# *Aeromonas*: Genomic Insights into an Environmental Pathogen and Reservoir of Antimicrobial Resistance

**DOI:** 10.1101/2025.05.22.655522

**Authors:** Nisha Singh, Rahma O. Golicha, Chetan Thakur, Mathew A Beale, Matthew J Dorman, Adrian Cazares, Alyce Taylor-Brown, Fatema-Tuz Johura, Mahamud ur Rashid, Shirajum Monira, Fatema Zohura, Tahmina Parveen, Sazzadul Islam Bhuiyan, Marzia Sultana, Balvinder Mohan, Daryl Domman, Christine Marie George, Samuel Kariuki, Munirul Alam, Neelam Taneja, Nicholas R Thomson

## Abstract

Aeromonads are an ecologically versatile group of bacteria that cause infection in aquatic animals and are recognised as an emerging human pathogen. Despite this, our understanding of *Aeromonas* diversity, especially the relationship between clinical and environmental strains, remains limited. Here, we present a complete view of the *Aeromonas* genus, comprising 1,853 genomes, and a detailed comparison of clinical and environmental strains from South Asia, including 996 newly sequenced genomes from Bangladesh and India. Phylogenetic analyses revealed that *Aeromonas* is a highly diverse genus, with no distinct clade separating clinical and environmental isolates. We identified 28 *Aeromonas* species and 905 novel sequence types, comprising 72.5% of the genomes. Notably, we show a high incidence of AMR genes across all isolates, including against front and last-line antibiotics. Finally, we highlight frequent misidentification of *Aeromonas* as *Vibrio cholerae*, key to cholera-endemic regions where both genera co-exist and are associated with diarrhoeal disease. Our study underscores *Aeromonas* as an important environmental AMR reservoir and emerging multi-species pathogen capable of spilling over into human populations.

## INTRODUCTION

Aeromonads (within the *Aeromonadaceae* family) are Gram-negative facultatively anaerobic bacteria with oxidase and catalase activity. *Aeromonas* spp. are abundant in aquatic habitats, with over 30 species identified^1^. At least 19 of these species are linked to human infections and are capable of causing significant diarrheal outbreaks. *Aeromonas* infections are common in children under 5 years old, especially in low- and middle-income countries (LMICs), where there is inadequate access to water and sanitation, resulting in high environmental exposure. The most common symptoms of *Aeromonas* infection include diarrhoea (which can be mistaken for cholera^2,3^), localised soft tissue infection, and bacteraemia^4–8^. Bacteraemia is most common in patients who have underlying health conditions, such as hepatobiliary disease, cancer, or diabetes^9^. There appear to be geographic differences in prevalence with <1% of cases of moderate to severe diarrhoea (MSD), attributable to *Aeromonas* in four African sites and Kolkata, India, whereas *Aeromonas* was implicated in >22% of MSD cases in children from Karachi, Pakistan, and Mirzapur in Bangladesh^10^. *Aeromonas* spp. are also important pathogens of animals, particularly fish and aquatic animals with significant economic consequences for aquaculture^11,12^.

Pathogenicity in Aeromonads is complex; they possess a variety of virulence factors that contribute to biofilm formation, cell adherence, invasion, and cytotoxicity. These include polar and lateral flagella^13,14^, adhesins^15^, lipopolysaccharides, iron-binding systems^16,17^, and numerous extracellular toxins and enzymes^18^ secreted by various systems such as type 2 and type 3 secretion systems^19,20^. Additionally, quorum-sensing systems play a crucial role in colonization and disease development^21–23^. Resistance to most classes of β-lactam antibiotics has also been observed among *Aeromonas* spp., but these bacteria have otherwise been reported to be susceptible to most first line antimicrobials, despite the widespread use of antibiotics in aquaculture^24^.

Notwithstanding the above, *Aeromonas* spp. are frequently misidentified as *Vibrio cholerae* in clinical and environmental samples. This is explained by sharing many biochemical and microbiological properties with other genera such as *Aerobacter, Pseudomonas, Escherichia, Proteus*, and *Vibrio*^3,25^. Both *Vibrio* and *Aeromonas* genera are oxidase-positive, able to thrive in aquatic and marine systems and exhibit similar colony morphologies on Thiosulphate Citrate Bile Salt Sucrose (TCBS) medium routinely used to select for *V. cholerae*^26^. These misidentifications pose serious epidemiological challenges, especially in regions where *V. cholerae* is endemic. Misidentification can not only lead to misreporting of cholera cases, but also a failure to implement necessary public health control measures. Hence, it is important to consider *Aeromonas* spp. in cholera surveillance and control efforts in endemic regions.

Recent genome data from 447 *Aeromonas* isolates from Pakistan revealed that human-associated *Aeromonas* in both cases and age-matched controls were highly diverse and displayed a high level of antimicrobial resistance (AMR), particularly extended-spectrum beta-lactamases (ESBLs)^27^. Looking into determinants that may explain the difference in clinical presentations, this study looked across an array of known or putative virulence factors, of which only two, *maf2* and *lafT* were weakly associated with the MSD cases. However, by focussing on isolates from humans, this study was unable to establish whether human pathogenic strains were a specialised subset of those found in the environment or if *Aeromonas* spp. infections are truly opportunistic linked to environmental exposure^27^.

Given the ubiquitous nature of Aeromonads in the environment, their potentially emerging role in MSD in South Asia, and the fact that they are often cultured from samples taken from patients with suspected cholera, we created a baseline population genomic snapshot of *Aeromonas* species from clinical and environmental samples. We sequenced two collections of *Aeromonas* isolates from *V. cholerae* endemic regions: Dhaka city, Bangladesh, and Northern India, together comprising 996 novel *Aeromonas* genomes. We compared their genetic diversity, predicted virulence gene profiles, and AMR profile, and contextualised this with genomes from a global *Aeromonas* population dataset.

## RESULTS

### Species diversity and distribution of *Aeromonas* spp. genomes

To examine the genomic diversity of *Aeromonas* strains across clinical and environmental settings in South Asia, we included largest and only available clinical genome datasets from Pakistan and environmental datasets from India and Bangladesh (this study). To provide a global context, we also incorporated genomes representing broad geographic and species diversity within the *Aeromonas* genus. In total, we assembled 1,853 *Aeromonas* genomes, including 996 newly sequenced genomes, of which 198 (10.68%) were mistakenly collected as *Vibrio cholerae* -133 from suspected cholera cases, households contacts in the ChoBI7 programme in Dhaka city^28^ and 65 from an environmental surveillance study for *V. cholerae* conducted in Bangladesh. Further, we generated 798 *Aeromonas* genomes from isolates taken from 274 water samples collected in Northern India. We contextualised our dataset using public archives, including 441 genomes of clinical origin from children with moderate to severe diarrhoea and corresponding healthy controls, taken in Pakistan^27^ as well as 416 globally distributed^29^ *Aeromonas* genomes from different sources. The largest proportion majority of publicly available genomes originated from Denmark (*n* = 100), followed by the United States (*n* = 71) and China (*n* = 62) (Supplementary Table 1).

We used GTDB-Tk (Genome Taxonomy Database) to perform species assignment, identifying 28 *Aeromonas* spp. across the 1,853 genomes, of which eight species dominated. *A. caviae* (*n* = 608) was the most numerous in our collection, followed by *A. veronii* (*n* = 566), *A. dhakensis* (*n* = 186), *A. salmonicida* (*n* = 176), *A. hydrophila* (*n* = 124), *A. enteropelogenes* (*n =* 105), *A. jandaei* (*n* = 25), *A. sanarellii* (*n* = 16), with the remaining 47 *Aeromonas* genomes representing 20 other *Aeromonas* species (Figure 1; summarised in Supplementary Figure 1a). To understand how our species predictions mapped to established species we used Average Nucleotide Identity (ANI) to evaluate their genetic relatedness. The intra-species ANI values ranged from 94.06% to 100%, falling slightly below the established definition for a single species (>95% ANI), in some instances. Conversely, inter-species ANI values ranged from 79.66% to 95.51% with higher-than-expected ANI and lower levels of expected divergence between *A. bestiarum* and *A. piscicola* (95.14-95.51%) (Supplementary Figure 2).

**Figure 1.**
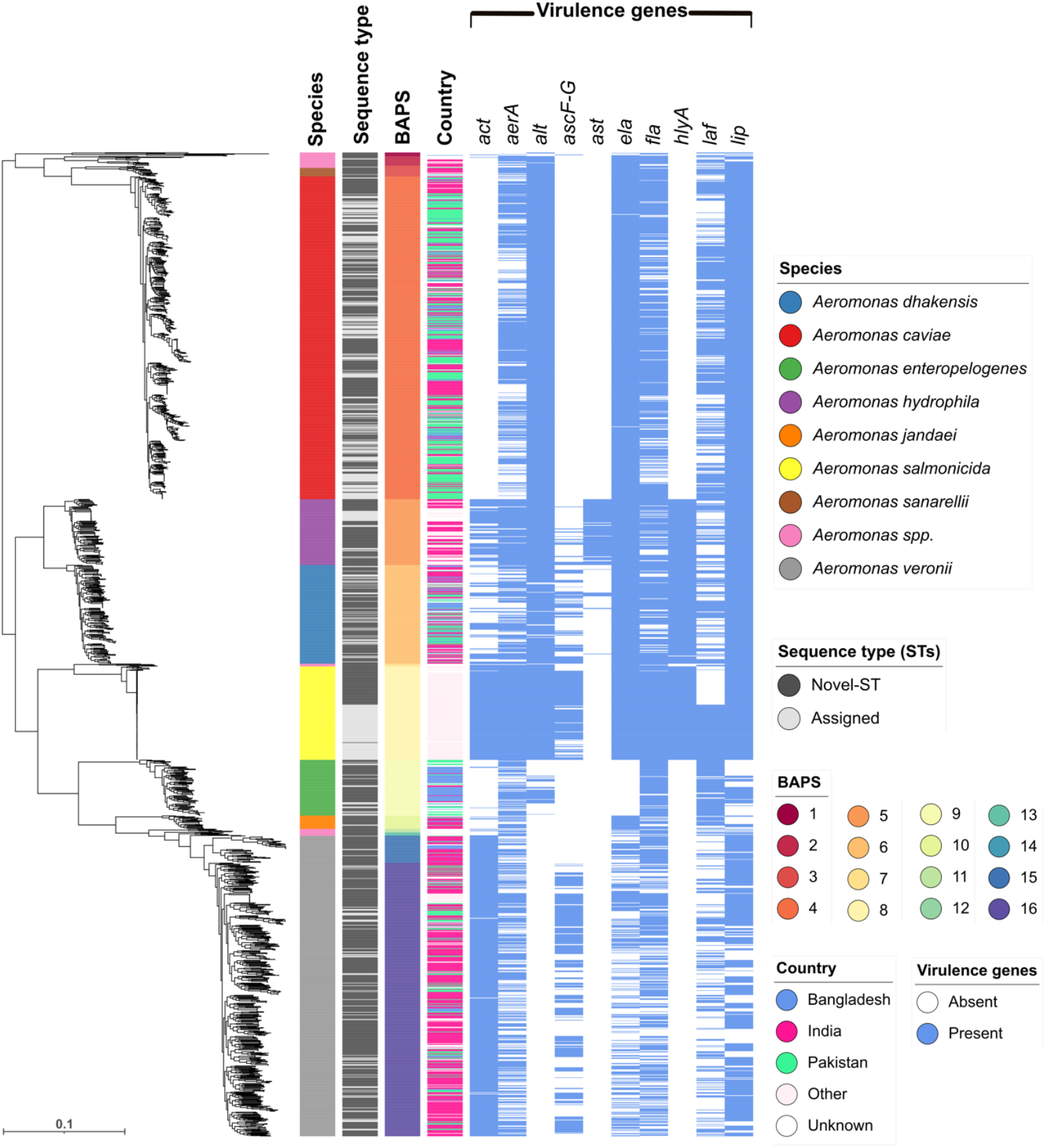
Taxonomic, genetic and geographic diversity of 1,853 *Aeromonas* species genomes. A maximum likelihood phylogenetic tree based on 1,969 core genes for *Aeromonas* spp. genomes. Refer to the key for interpretation of the colour strips alongside. The scale indicates an evolutionary distance of 0.1 nucleotide substitutions per site. BAPS refers to Bayesian Analysis of Population Structure, used here to delineate genetic clusters.

Looking at the geographic distribution of *Aeromonas* species, across both clinical and environmental samples from India, Pakistan and Bangladesh where we had the densest sampling (1,438 *Aeromonas* genomes; Supplementary Figure 1b). *Aeromonas* spp. found across all three South Asian countries included *A. caviae, A. dhakensis, A. enteropelogenes, A. jandaei, A. veronii* and *A. sanarellii*. Conversely, *A. sanarellii* and *A. jandaei* were rare in all three countries, whilst overall there were similarities in the relative abundance of the different species, despite differences in sampling depth and strategy (see methods), there were also clear differences: *A. caviae* predominated in Bangladesh and Pakistan (31.81% and 64.39%, respectively) and *A. veronii* dominated the samples collected in India (48.31% of isolates). Other species, such as *A. schubertii*, were exclusively seen in Bangladesh (*n* = 1), while *A. media* (*n* = 4) and *A. rivipollensis* (*n* = 5) were exclusively seen in India (Supplementary Figure 1b). What is more, environmental samples from Northern India could include between 1-5 different *Aeromonas* species (Supplementary Figure 3a). For countries where we had less dense sampling, *A. hydrophila* was the most common in the United States (42.25%; 30/71), whereas *A. salmonicida* was the dominant species in the samples from Denmark (99%, 99/100) and China (96.77%; 60/62).

In India, all samples were collected as part of a cholera surveillance study. Samples were taken from known cholera hotspots and at times of the year when cholera cases were more common. All water samples were cultured on *V. cholerae* selective media (TCBS; see methods). Using MALDI-TOF it was possible to look at the *Aeromonas* spp. positivity rate, as well as for the presence of a range of other enteric pathogens (Supplementary Figure 4). Out of 408 collected water samples from Northern India, 274 samples (67.15%) tested positive for *Aeromonas* spp., 145 (35.53%) for *V. cholerae*, 112 (27.45%) for *E. coli*, 198 (48.52%) for *Klebsiella* spp. and 35 (8.57%) for non-cholera *Vibrio* spp. (8.57%, 35/408; including species *V. fluvialis, V. navarrensis, V. cidicii, V. injensis* and *V. metschnikovii*).

Of the samples positive for *Aeromonas* spp., *V. cholerae* was also isolated from 91 samples (33.57%, 91/274). These co-occurrences were primarily found in river samples (75/91), followed by ponds (13/91) and canals (4/91), across different regions of Northern India. A total of seven *Aeromonas* species: *A. veronii, A. caviae, A. hydrophila, A. dhakensis, A. jandaei, A. enteropelogenes* and *A. sanarellii*–were identified as cohabiting with *V. cholerae*. However, there was no clear association between the co-occurrence of *V. cholerae* with any one specific *Aeromonas* species, these data largely mirrored the relative isolation rates for *Aeromonas* species from Northern India (Supplementary Figure 3b; Supplementary Figure 3c).

### Genomic Taxonomy of *Aeromonas* spp

To infer the phylogeny of the genus, we standardised the annotation of our underlying dataset using PROKKA and then defined the *Aeromonas* spp. pangenome using Panaroo. This comprised 47,542 coding sequences (CDS), of which 1,969 were defined as core (found in more than 99% of genomes) and 118 as soft core (found in 95-99% of genomes), with 45,455 accessory genes (found in 0-95% of genomes). We constructed a maximum likelihood phylogenetic tree from a core gene alignment (Figure 1; see methods). We used Bayesian Analysis of Population Structure (BAPS, implemented in fastBAPS) to delineate genomic clusters and correlated these with ANI values and the species assignments from GTDB-Tk. From BAPS, whilst most taxa fell on deeply branching nodes in the *Aeromonas* spp. tree with high concordance with the ANI defined species blocks (Figure 1; Supplementary Figure 2), *A. veronii* fell across two BAPS clusters (15 and 16). Conversely, BAPS clusters 1, 2, 3, and 7 each span multiple species of *Aeromonas* (Supplementary Table 2). These inconsistencies may reflect differences between the way the BAPS model clusters genomes, which, unlike ANI, is not distance based.

Next, to determine how much of the diversity within known taxonomic groups had been previously observed, we constructed *in silico* MLST^30^ profiles. Of the 1,853 MLST profiles generated, only 495 (26.71%) genomes were assigned to 224 known MLST allelic profiles, with 1,343 genomes representing 905 novel sequence types (STs). After assignment by pubmlst.org, 1,838 of our *Aeromonas* spp. genomes were assigned to 1,129 STs (15 genomes remained unassigned; see Supplementary Information-I), now accessible through the *Aeromonas* MLST database^31^. Of note these novel STs define discrete branches within all species, while 836 STs represented singletons, with clusters of genomes belonging to the same ST being rare. The exceptions to this were ST-2 (*n* = 103) and the newly defined ST-1799 (*n* = 60) for *A. salmonicida*, as well as ST-2398 (*n* = 20) and ST-251 (*n* = 19), representing *A. caviae* and *A. hydrophila* genomes respectively.

### The relationship between clinically and environmentally derived *Aeromonas* spp. isolates and human health

Several virulence genes have been previously described amongst *Aeromonas* species, including the haemolytic toxins: aerolysin-related cytotoxic enterotoxin (*act*), heat-labile cytotonic enterotoxin (*alt*), heat-stable cytotonic toxins (*ast*), hemolysin (*hlyA*), cytotoxin aerolysin (*aerA*) as well as polar flagellum (*fla*), lateral flagella (*laf*), the zinc-and-iron-dependant metallo-endopeptidase elastase (*ela*), the lipase (*lip*) and the type III secretion system (TTSS) encoded by *ascF-G*. To determine whether any of the known virulence genes are linked to the particular *Aeromonas* species, we used *in silico* PCR to understand the distribution of these genes across 1,853 *Aeromonas* species genomes.

We analysed patterns of virulence gene distribution using principal component analysis (PCA) and hierarchical clustering (Figure 2a; Figure 2b). The PCA revealed that certain species, such as *A. hydrophila, A. dhakensis, A. salmonicida*, and *A. veronii*, formed relatively well-defined clusters suggesting distinct virulence profiles, with some overlap between the phylogenetically related *A. hydrophila* and *A. dhakensis*, as well as between the phylogenetically distant *A. hydrophila* and *A. salmonicida* (Figure 1; Figure 2a). Genes such as *aerA, fla, laf*, and *lip* were present across all species (e.g. *aerA* and *fla* were present in >50% of genomes tested from all of taxa screened), whilst others showed deep associations with a particular species, consistent with previous studies^32,33^. *A. caviae* (*n* = 608) and *A. salmonicida* (*n* = 176) carried the highest proportion of some virulence genes, with all genomes of these species possessing *alt, ela* and *lip* genes, whilst *ela* and *lip* were also uniformly present in *A. dhakensis, A. hydrophila*, and *A. sanarellii* genomes (Figure 1; summarised in Figure 3a).

The Type III secretion system genes (*ascF-G*) were found more often in *A. salmonicida* (141/176*), A. veronii* (288/566), *A. dhakensis* (78/186), and *A. hydrophila* (21/124). Gene *hlyA* was absent in *A. jandaei, A. enteropelogenes, A. sanarellii, A. caviae*, and *A. veronii*, representing a potential molecular marker for differentiating between species (Figure 2b). Similarly, *ast* gene, which encodes an enterotoxin, was found only in *A. hydrophila* (*n* = 118) and *A. dhakensis* (*n* = 11). We noted, a high prevalence of the combination of the cytotoxic enterotoxin (*act*) and cytotonic heat-labile enterotoxin (*alt*) in *A. salmonicida* (*alt* = 100%, *act* = 99%), *A. hydrophila* (*alt* = 100%, *act* = 72%), and *A. dhakensis* (*alt* = 94%, *act* = 29%), and the distribution *laf*, suggesting this gene had been gained or lost multiple times across the phylogeny (Figure 1): for example it was present in *A. salmonicida* ST-2 but absent from ST-1799.

**Figure 2.**
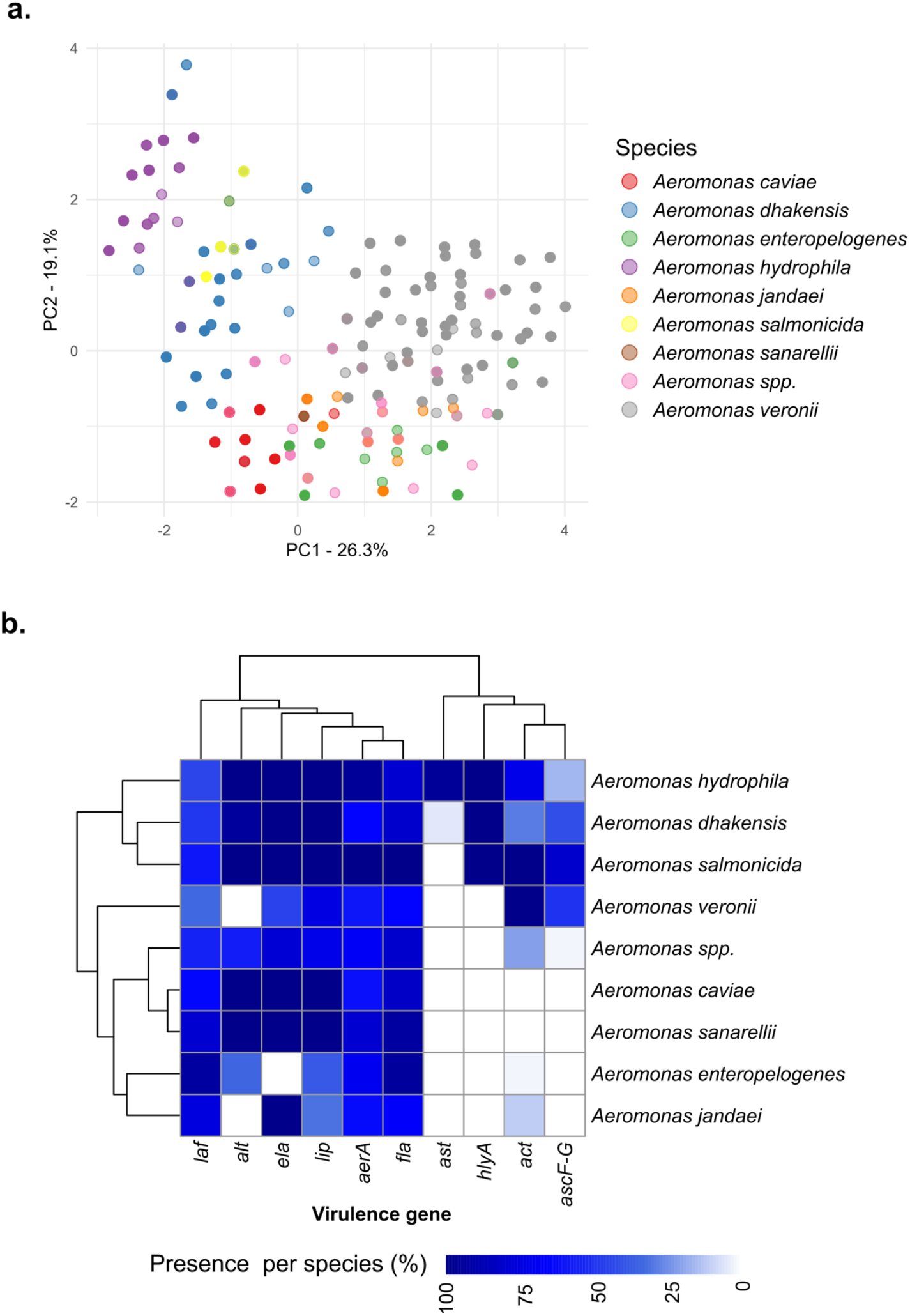
Distribution of virulence genes across 1,853 *Aeromonas* species genomes. **(a)** Principal Component Analysis (PCA) of virulence genes across different *Aeromonas* spp. (see key for coloured dots representing species). **(b)** Hierarchical clustering of virulence genes across *Aeromonas* spp., with colours representing the percentage presence of genes (see key for scale).

Lastly, we investigated whether there were associations between species and sample origin. For this we focussed on South Asia (Bangladesh, India and Pakistan), since here we had 1,438 *Aeromonas* spp. genomes confirmed to originate from clinical or environmental samples. As shown in Supplementary Figure 5a and 5b, there was no obvious association between species and source. This was also true when looking at *A. caviae* taken from South Asia (*n* = 577), where we had almost equal representation from clinical (*n* = 286; of which 153 were from individuals with moderate to severe diarrhoea, while 133 were asymptomatic) and environmental (*n* = 291) *Aeromonas* genomes (Supplementary Figure 6). Of the 12 BAPS clusters that contained both clinical and environmental *Aeromonas* genomes, the only exception was cluster-4.11, which included just five isolates in total, all from environmental sources (Supplementary Table 3).

### *In silico* prediction of AMR

To understand if there were differences in AMR profiles across or within species, we inferred the genotypic AMR profiles for all the genomes included in this study. Overall, we found 162 discrete AMR genes, conferring resistance to 16 different antimicrobial classes including: β-lactams, sulphonamides, tetracycline, and aminoglycosides (Figure 3b, summarised in Supplementary Table 4). Of these, those conferring resistance to β-lactam antibiotics were the most common (40.74%, 66/162), including chromosomally encoded metallo-β- lactamases, cephalosporinases, and oxacillinases (Supplementary Table 5). However, AMR genes *mcr-3, mcr-3*.*12, mcr-3*.*3, mcr-3*.*6, mcr-3*.*8, mcr-5*, and *mcr-7*.*1*, which confer resistance to colistin (often considered as last-resort of antimicrobials), were detected across eight species, including *A. caviae, A. hydrophila, A. jandaei, A. media, A. piscicola, A. salmonicida, A. sobria* and *A. veronii* (Supplementary Table 4; see Supplementary Information-I). Only three genomes in our collection lacked any known AMR gene.

**Figure 3.**
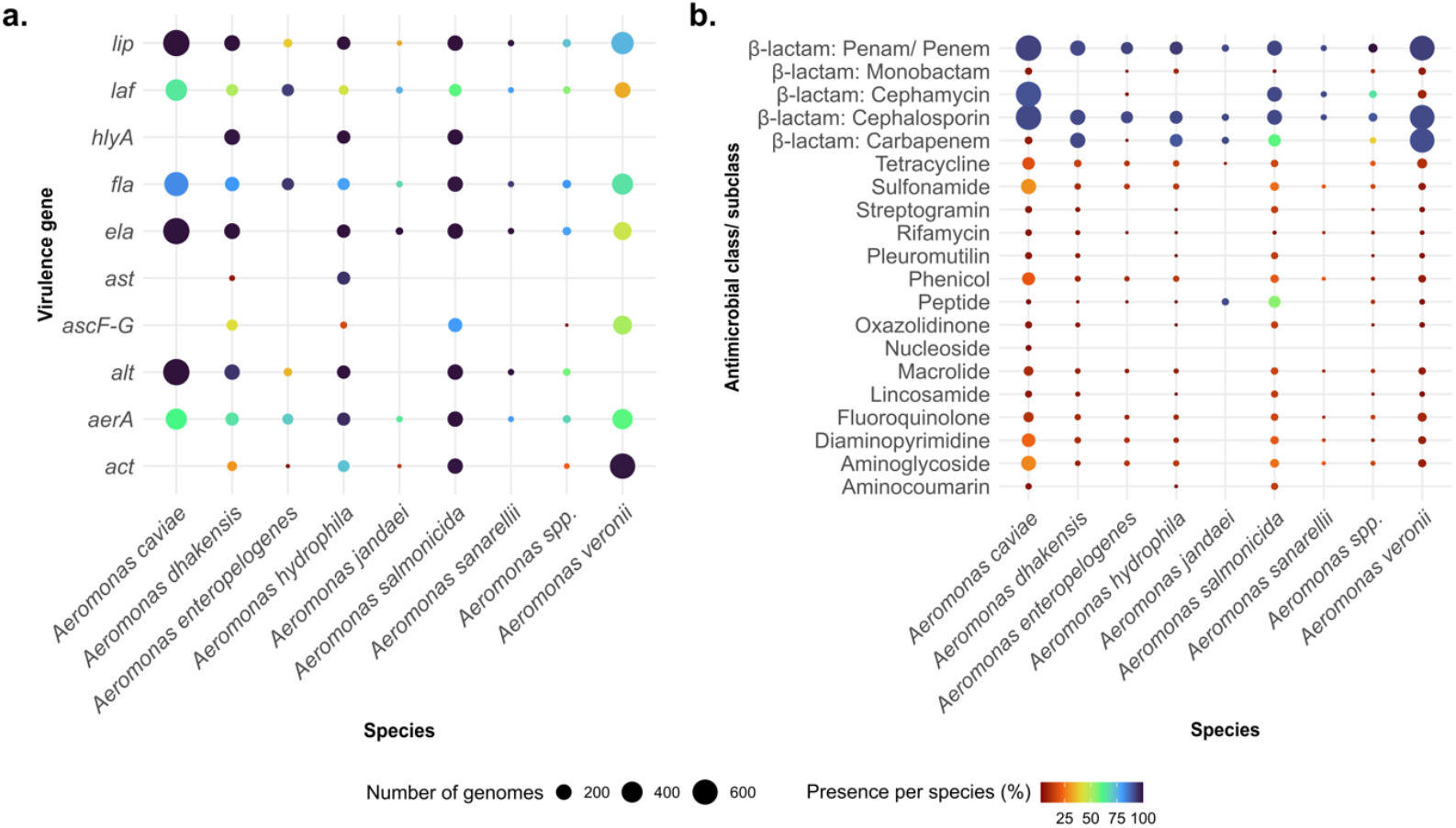
Distribution of virulence genes and drug classes corresponding to antimicrobial resistance across 1,853 *Aeromonas* species genomes. The X-axis represents the most abundant *Aeromonas* species, with less abundant species grouped as *Aeromonas* spp., and the Y-axis denotes virulence genes in **(a)** and antimicrobial classes corresponding to AMR genes in **(b)**. Each dot is coloured based on the gene abundance per species (see scale), and the size of the dots corresponds to the number of genomes.

Looking more closely at the subset of 1,438 South Asian genomes, we found AMR patterns varied significantly across India, Bangladesh, and Pakistan (Figure 4a). *Aeromonas* isolates from Pakistan exhibited the highest prevalence of resistance to β-lactams (cephamycin: 65.6%), sulfonamides (24.3%), diaminopyrimidines (22.4%), aminoglycosides (22.2%), tetracyclines (18.4%), phenicols (16.3%), and fluoroquinolones (12.9%). Indian isolates showed comparatively lower prevalence of resistance gene presence amongst the three South Asian countries, with the highest resistance observed for carbapenems (69.2%); macrolides (7.8%); rifamycins (3.3%); and oxazolidinones, pleuromutilins, and streptogramins (4.4%). *Aeromonas* from Bangladesh had the lowest prevalence of resistance, with no genes conferring resistance to lincosamides, oxazolidinones, pleuromutilins, streptogramins, and nucleosides. Notably, all genomes across the three countries were predicted to carry cephalosporin resistance genes. Among these, 4.5% of the isolates (65/1,438; 14 clinical and 51 environmental) carried genes encoding third-generation extended-spectrum β-lactamases (ESBLs), such as *bla*_CTX-M_, *bla*_GES_, *bla*_PER_, *bla*_TEM_, and *bla*_VEB_, highlighting the problem of widespread β-lactam resistance, consistent with global trends^27,34,35^. This is particularly concerning given the lack of standardized treatment guidelines for *Aeromonas* infections, where fluoroquinolones, particularly ciprofloxacin and, in some cases, third-generation cephalosporins are commonly used empirically ^25,36–39^.

**Figure 4.**
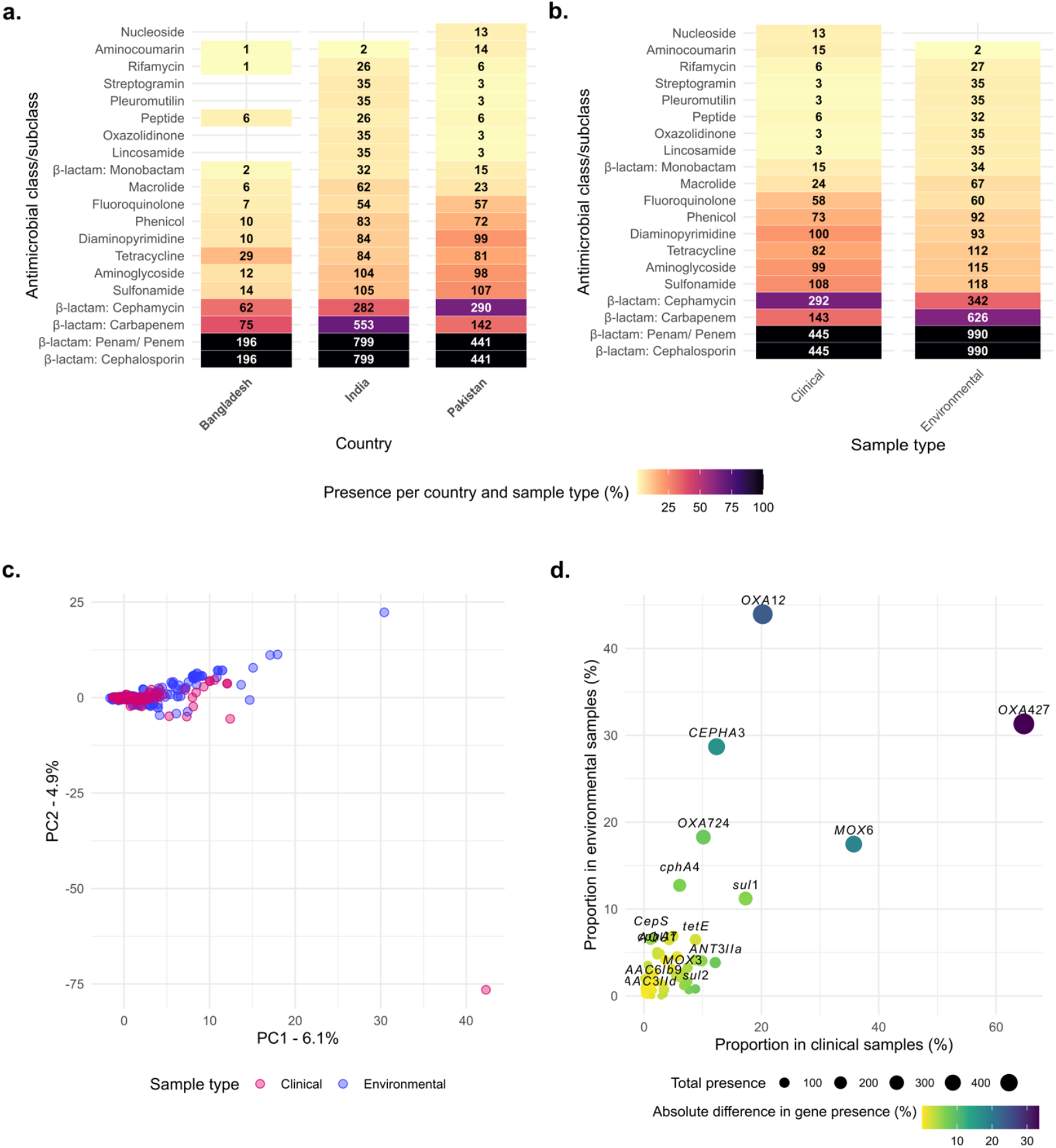
Distribution of drug classes corresponding to antimicrobial resistance genes across 1,436 *Aeromonas* species genomes in South Asia, including Bangladesh, India, and Pakistan. The X-axis represents the country of origin in **(a)** and the sample type (clinical vs. environmental) in **(b)**, while the Y-axis denotes antimicrobial classes corresponding to resistance genes. The heatmap is color-coded to indicate the percentage presence of AMR genes per country and sample type, with numeric indicators above each tile denoting the number of genomes carrying the gene within that category. **(c)** Principal Component Analysis (PCA) illustrates the clustering of genes in clinical and environmental isolates (see key for coloured dots). **(d)** Absolute difference in gene presence between clinical and environmental isolates, with yellow-green indicating minimal or no difference and blue-violet representing genes with significant variation in presence across sample types.

To assess the genetic differences in AMR profile between clinical and environmental isolates we used PCA, which revealed substantial overlap between both groups, indicating similar gene profiles overall (Figure 4c). However, certain genes exhibited notable differences in prevalence between the two groups (Figure 4d). Clinical isolates had a higher proportion of *bla*_OXA-427_ (64.7%), *bla*_MOX-6_ (35.7%), and *sul1* (17.3%) compared to environmentally derived isolate genomes. In contrast, environmental isolates showed higher proportions of *bla*_OXA-12_ (43.9%), *bla*_OXA-724_ (18.3%), *bla*_CEPHA3_ (28.7%) and *bla*_*c*phA4_ (12.7%), Figure 4d. Of note, one Indian environmentally derived *A. sanarellii* isolate was predicted to be multidrug resistant by possessing genes to seven drug classes including: aminoglycoside, β-lactam, diaminopyrimidine, macrolide, phenicol, rifamycin, and sulphonamide (see Supplementary Information-I).

## DISCUSSION

We initiated this study to provide a genomic framework for understanding the relationship between *Aeromonas* spp. from the natural aquatic environment and those isolated from clinical samples linked to disease. We also wanted to understand the nature and diversity of *Aeromonas* spp. co-inhabiting the same natural environment with *Vibrio cholerae*, in endemic settings. We included a range of isolates and genomes generated here, as well as published Aeromonads genomes from moderate to severe diarrhoea cases and matched controls from Pakistan^27^. We investigated the genetic diversity, prevalence of virulence and antibiotic resistance genes amongst Aeromonads genomes from households and freshwater in multiple countries. In doing so, we have provided the most comprehensive view of this genus thus far.

Among 1,853 genomes included in this study, we identified 28 *Aeromonas* species, including *A. veronii, A. caviae, A. dhakensis, A. enteropelogenes, A. salmonicida*, and *A. hydrophila*, several of which are known human or fish pathogens^37^. Notably, *A. dhakensis* is overrepresented in our study, constituting ∼22% of all isolates from Bangladesh. Other notable geographic signals include the psychrophilic *A. salmonicida*, seen largely in cooler countries such as Denmark (56%) and certain regions of China (35%), where most commercial fish farming is done^40^. Here, the majority of the *A. salmonicida* were collected from infections of trout, *Oncorhynchus mykiss*, known to be highly susceptible to *A. salmonicida* infections. Similarly, 20% of *A. hydrophila* described here were from the United States of America, where *A. hydrophila* causes significant economic losses in catfish farming^41^.

Despite being considered an emerging human and animal pathogen linked to a wide array of diseases, there is limited available genomic data for the *Aeromonas* genus, particularly from environmental sources. Our core gene phylogeny highlights that this genus is remarkably diverse; this was particularly evident from the high number of novel MLST STs (905), with 1,343 (72.47%) genomes assigned to these previously unreported profiles (see Supplementary Information-I). Perhaps unsurprisingly given the extent of the previously unknown diversity, ANI showed the intra-species ANI values for *A. sobria* and *A. veronii* fell below the standard 95% threshold, challenging current boundaries for species delineation and suggesting a need for reclassification within these species.

Looking at the species delineation between human and environmental samples, from previous studies of children with moderate-to-severe diarrhoea in Karachi, Pakistan, *A. caviae* (64.2%) was seen to be the most numerous, followed by *A. veronii* (19.2%), *A. dhakensis* (9.8%), and *A. enteropelogenes* (4.9%)^27^. Other studies have also shown that *A. hydrophila* is also common in disease cases^42^. From our data, comparing clinical and environmentally derived isolates from South Asia, we showed a similar distribution across both human and environmental samples, with *A. caviae, A. veronii, A. dhakensis*, and *A. enteropelogenes* being the most abundant.

To identify signatures of intra- and interspecific ecological separation within and between clinical and environmental isolates, we examined all genomes for differences in virulence potential and AMR profiles, linked to taxonomic affiliation and source of isolation. From the virulence gene profiles, there were no significant genetic differences between clinical and environmental strains. However, these profiles allowed us to observe distinct clusters formed by species such as *A. hydrophila, A. veronii, A. dhakensis*, and *A. salmonicida*, implying that virulence gene profiles could be used to classify *Aeromonas* spp. into different groups.

Most *Aeromonas* genomes, regardless of source, carried AMR genes, particularly those conferring resistance to β-lactams, aminoglycosides, and sulphonamides, consistent with previous clinical studies^27,40^. Notably, we identified a wide range of β-lactamase genes, including chromosomally encoded class B metallo-β-lactamases (*bla*_CphA_), class C cephalosporinases (*AmpC* β-lactamases: *bla*_FOX_, *bla*_MOX_), and class D β-lactamases (oxacillinases: *bla*_OXA_ variants), underscoring the well-documented intrinsic resistance mechanisms of *Aeromonas* spp ^43–45^. Additionally, a substantial number carried acquired ESBLs (e.g., *bla*_CTX-M_, *bla*_GES_, *bla*_PER_, and *bla*_VEB_) and carbapenemases (*bla*_KPC_, *bla*_VIM_)^46,47^. Our analysis showed considerable overlap in AMR profiles between clinical and environmental isolates, with few genes showing distinct patterns. For example, *bla*_OXA-427_, a class D carbapenemases previously identified in several *Enterobacteriaceae* clinical strains^48^, appeared more frequently in clinical *Aeromonas* isolates. In contrast, The class D carbapenemases gene *bla*_*O*XA-12_, first identified in *A. jandaei* ^49^ and most abundant in environmental isolates, was found in all *A. jandaei*, 565 *A. caviae*, and two *A. hydrophila* isolates, as well as in less common species like *A. allosaccharophila* (*n* = 5), *A. australiensis* (*n* = 1), *A. fluvialis* (*n* = 1), and *A. sobria* (*n* = 6). The gene *bla*_*O*XA-12_ was widely distributed across several countries and has been previously reported in environmental *A. veronii* from the United States^50^ and in strains from humans and animals in India^51^. The presence of these genes in both clinical and environmental isolates raise concern, as carbapenems are critical last-resort antibiotics^52,53^.

We also observed the presence of mobile colistin resistance genes in eight different *Aeromonas* species, spanning diverse geographical regions including Bangladesh, China, Denmark, India, Pakistan, Spain, the United Kingdom, and the United States. Colistin remains a last-resort antibiotic for treating clinical infections caused by multidrug-resistant Gram-negative bacteria^54^. The presence of *mcr* genes, many of which are plasmid-encoded, raises significant public health concerns, as they can facilitate horizontal gene transfer to other clinically important pathogens^54^. The widespread occurrence of *mcr* in both clinical and environmental *Aeromonas*^55–57^ isolates further suggests that this genus may act as both a reservoir and conduit for global colistin resistance dissemination.

Notably, the AMR patterns observed in *Aeromonas* spp. from South Asia highlight significant regional variability, with Pakistan^27^ exhibiting the highest levels of resistance gene carriage across multiple antibiotic classes. This may be attributed to all Pakistani isolates in our study originating from clinical samples taken from individuals with moderate to severe diarrhoea or corresponding controls.

Taken together, the widespread occurrence of clinically relevant resistance genes in *Aeromonas*, a genus found in aquatic and wastewater environments, highlights its role as a key environmental sentinel in the One Health context. This aligns with the broader understanding that various environmental sources contaminated with residual antimicrobial substances, such as wastewater, agricultural runoff, and pharmaceutical discharges^58–60^, select for AMR in important bacteria such *Aeromonas* and members of the *Enterobacteriaceae*. These contaminated environments not only support the survival of resistant organisms but also promote their genetic exchange and adaptation, potentially accelerating the spread of resistance across both clinical and environmental settings and harbouring genes conferring resistance to first and last line antimicrobials.

One of the drivers for this study was that *Aeromonas* spp. are frequently misidentified as *V. cholerae* in clinical and environmental samples^3,61^. Our data shows that 67.15% of the environmental isolates cultured on TCBS in Northern India were *Aeromonas* spp. and not *V. cholerae*. Similarly, of the environmental isolates that were originally collected and identified as *V. cholerae* through culture from the Dhaka household study, ∼70% were subsequently reclassified as *Aeromonas* spp. This contrasts the prevalence of this genus among the 239 stool samples positive for *V. cholerae* from the same study, where only two *Aeromonas* spp. were misidentified as *V. cholerae*, both of which were cultured from asymptomatic individuals (^28,^ this study; Dr Munir Alam personal communication). This suggests that misidentification of *Aeromonas* for *V. cholerae* is unlikely to overestimate cholera prevalence in clinical settings (e.g. from human stool). Still, environmental sampling and identification based on microbiological growth and appearance on selective media may overinflate apparent *V. cholerae* prevalence in the environment, especially during a cholera epidemic, if culture is the only confirmatory method employed.

In conclusion, our data brings together the different ecological views of this bacterium, it shows the environment to be an important reservoir of a highly diverse range of species capable of overspilling into humans and in some instances causing disease. It is also clear that they carry an important array of therapeutically relevant AMR genes. Here, by taking a broader approach to understanding questions such as sources and sinks of AMR and considering non-traditional organisms, we show that *Aeromonas* has seemingly been hidden in plain sight – both as an emerging human pathogen and clearly as an important environmental reservoir of AMR.

## METHODS

### *Aeromonas* isolates and genome sequences

We used a collection of 996 novel *Aeromonas* isolates. Of these, 198 had been originally cultured and identified as *V. cholerae* using conventional microbiological and biochemical techniques^62^ before they were sequenced and classified genomically as *Aeromonas*. These included 133 isolates from a study entitled the Cholera-Hospital-Based-Intervention-for-7-Days (ChoBI7) programme in Dhaka city^28^ and 65 isolates from an environmental surveillance study for *V. cholerae* conducted in Bangladesh. The remaining 798 genomes were obtained from 408 water samples collected for isolation of *V. cholerae* from ponds, rivers, and canals across six different states - Delhi, Haryana, Himachal Pradesh, Punjab, Uttar Pradesh and Uttarakhand as well as Chandigarh in Northern India - between 2020 and 2023. The collected water samples were first enriched in alkaline peptone water and Selenite F broth. They were then sub-cultured onto various media. Blood agar and Thiosulphate Citrate Bile Salt Sucrose (TCBS; Difco Laboratories, Detroit, Michigan, USA) agar were used for the isolation of *V. cholerae* and *Aeromonas* spp. MacConkey agar (HiMedia Laboratories Private Limited, Mumbai, India) and Xylose Lysine Deoxycholate (XLD; Difco Laboratories, Detroit, Michigan, USA) were used for isolating other *Enterobacteriaceae*. The presumptively selected colonies from media listed above were identified by matrix-assisted laser desorption ionization-time of flight mass spectrometry (MALDI-TOF)^63^ and isolates identified as *Aeromonas* spp. underwent DNA extraction and whole genome sequencing as described below. Between 1 and 9 colonies were retained from each *Aeromonas*-positive water sample in India for downstream processing, including DNA extraction and whole genome sequencing.

DNA was extracted from bacterial isolates using the Wizard® Genomic DNA Purification Kit (Promega, Madison, WI, USA) and Qiagen DNeasy Blood & Tissue Kits (Qiagen, Hilden, Germany). DNA extracts were subjected to a standardised library preparation and submitted for whole genome sequencing (WGS) on the HiSeq X10 platform (Illumina, San Diego) at the Wellcome Sanger Institute (Cambridgeshire, UK).

Genomes were contextualised using 416 *Aeromonas* genomes (including one genome from India) downloaded from a library of 661,405 previously assembled bacterial genomes held in the European Nucleotide Archive (ENA)^29^, along with 441 genomes from a study on children with moderate to severe diarrhoea and corresponding controls in Pakistan^27^.

### Quality control and genome assembly

We performed initial quality control of sequencing reads using FastQC (v0.11.4.)^64^ and MultiQC (v1.17)^65^, removing read sets that demonstrated bimodal peaks in GC content or substantially abnormal read quality metrics. We then performed *de novo* assembly using SPAdes (v3.9.0)^66^ to obtain contiguous scaffolded assemblies. Additional quality control was performed on the *de novo* assemblies using Quast (v5.0.2)^67^ and CheckM (v1.1.2)^68^ to calculate the genome contamination and completeness. Assemblies were filtered to retain high-quality assemblies based on genome length (4.5-5.2 MB), genome contamination (<5%) and genome completeness (>95%). Overall, 5% (97/1,950) of the genomes were excluded from the study based on these criteria, including six genomes from the Pakistani study that failed our quality control.

### Species identification and distinction

We initially used Kraken2 (v2.2)^69^ and Bracken (v2.6.2)^70^ (Standard RefSeq database containing archaea, bacteria, viruses, plasmids and human, downloaded from https://benlangmead.github.io/aws-indexes/k2 on 9th October 2023) to assign taxonomic labels and assess the species abundance, respectively. Additionally, GTDB-Tk (v. 2.3.0)^71^ was used alongside Kraken2 and Bracken to enhance taxonomic classification of assemblies. This was followed by assessment of genome-relatedness indices, where we used FastANI (v1.3)^72^ for the pairwise comparison of the *Aeromonas* genomes. The FastANI threshold was set at 90% based on pairwise sequence mapping ^72^, and heatmaps based on estimated ANI for all *Aeromonas* genomes were created using the Complex heatmap package in R (v4.4.3)^73^.

### Prediction of AMR and virulence genes

AMR genes were identified from the assembled contigs using ABRicate (v1.0.1, available at https://github.com/tseemann/abricate) to query the Comprehensive Antimicrobial Resistance Database (CARD, v3.2.4)^74^ where the CARD identification threshold was set at 80% of nucleotide identity. For the identification of the putative virulence genes, we initially used ABRicate with the Virulence Finder Database^75^, but this did not yield any hits. We subsequently used in_silico_pcr (v1.0.0, available at https://github.com/sanger-pathogens/sh16_scripts/blob/master/legacy/in_silico_pcr.py) with previously described primer sequences (see Supplementary Table 6)^76,77^ to detect virulence genes. To explore the patterns of AMR and virulence gene distribution in relation to sample origin (clinical vs. environmental) and *Aeromonas* species, we conducted a Principal Component Analysis (PCA) using the prcomp function from the stats package in R (v4.4.3)^73^, based on presence/absence matrices. The first two components, PCA1 and PCA2, were selected and visualized using the ggplot2 package.

### Genome annotation and phylogenetic analysis

Draft *Aeromonas* genomes were annotated using PROKKA (v1.4.5)^78^ with default parameters. We used panaroo (v1.3.0)^79^ to infer the pangenome of 1,853 genomes and used the resulting core gene alignment (comprising 1,969 core genes) for phylogenetic analysis. We extracted 1,282,613 variable sites using SNP-sites (v2.51)^80^ and the resulting multiple sequence alignment file was used to infer a phylogeny with IQ-TREE (v2.1)^81^, using a generalized time reversible substitution model with a FreeRate model of heterogeneity^82^, incorporating invariant sites using the ‘-fconst’ parameter, and with 1,000 UltraFast Bootstraps^83^. Phylogenetic trees and associated metadata were visualized in iTOL (v.5.6)^84^. Similarly, a separate phylogenetic tree was constructed for genomes from India, Bangladesh, and Pakistan (*n* = 1,438), consisting of 2,067 core genes and 1,197,118 variable sites.

### Multi-locus Sequence Typing (MLST) and Subtyping of *Aeromonas* isolates

We used the MLST tool (v2.9, available at https://github.com/tseemann/mlst) to infer MLST against the *Aeromonas* scheme developed by Martino and colleagues^30^ and hosted at https://pubmlst.org/bigsdb^85^. This MLST scheme uses six housekeeping genes (*gyrB, groL, gltA, metG, ppsA*, and *recA*) to subtype *Aeromonas* below the species level into Sequence Types (STs). Novel alleles and STs were submitted to the *Aeromonas* MLST database via pubmlst.org for naming.

Since individual genomes were distributed across many different MLSTs, we also employed Bayesian Analysis of Population Structure (BAPS) using fastBAPS (v1.0.6)^86^ to define higher level grouping within the species.

## DATA AVAILABILITY

Short-read sequence data (150 bp paired-end reads) are available in the EMBL Nucleotide Sequence Database (European Nucleotide Archive, http://www.ebi.ac.uk/ena). For detail of accession number, see the Supplementary Information-I. Additional data supporting the findings of this study are available in the Supplementary Data.

## ACKNOWLEDGEMENTS

This research was funded in whole, or in part, by the Wellcome Trust (220540/Z/20/A and 215704/Z/19/Z). For the purpose of open access, the authors have applied a CC-BY public copyright licence to any author-accepted manuscript version arising from this submission. The authors thank the teams involved in sample collection in Bangladesh and India, and the Parasites and Microbes Programme Sequencing and Informatics teams at the Wellcome Sanger Institute for support. We would like to thank the National Biodiversity Authority of India for their support. MA acknowledges with gratitude the contribution of all collaborating colleagues of the grant in Bangladesh and the USA, including the lab and field team members of icddr,b. icddr,b acknowledges the donors providing unrestricted support for its operation and research.

## AUTHORS CONTRIBUTION

Conceptualization: NS, RO, MAB, MCD, AC, ATB, NRT Formal analysis: NS, RO, MAB, NRT

Investigation: NS, CT, FTJ, MR, SM, FZ, TP, SIB, MS, MA, NT, NRT

Resources: MA, NT, NRT

Writing and Visualization: NS, RO, MAB, NRT Supervision: MAB, MCD, AC, ATB, SK, NT, MA, NRT

Review & Editing: NS, RO, CT, MAB, MCD, AC, ATB, FTJ, MR, SM, FZ, TP, SIB, MS, BM, DD, CMG, SK,

MA, NT, NRT. All authors reviewed the manuscript.

