## Supplemental Material for "*Aeromonas*: Genomic Insights into an Environmental Pathogen and Reservoir of Antimicrobial Resistance"

### SUPPLEMENTARY DATA (FIGURES)

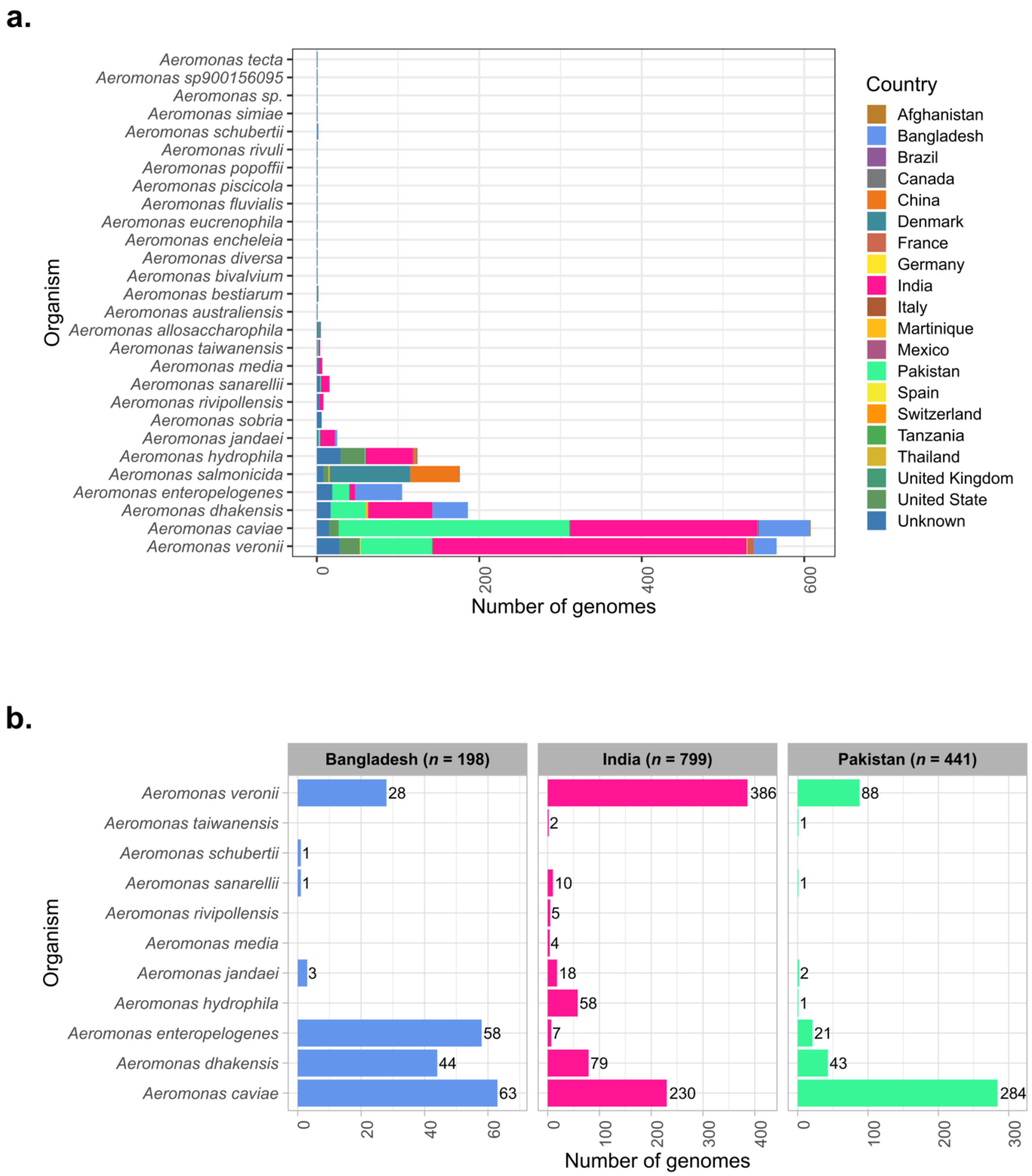


**Supplementary Figure 1**: **Geographic distribution of *Aeromonas* species genomes included in this study (a)** A total of 1,853 *Aeromonas* species genomes, including 996 sequenced in this study. Coloured bars represent various *Aeromonas* species across different countries (see key). **(b)** A subset of 1,438 *Aeromonas* species genomes distributed across South Asia, including Bangladesh, India, and Pakistan. The number in front of each bar indicates the number of genomes for the respective species in each country.

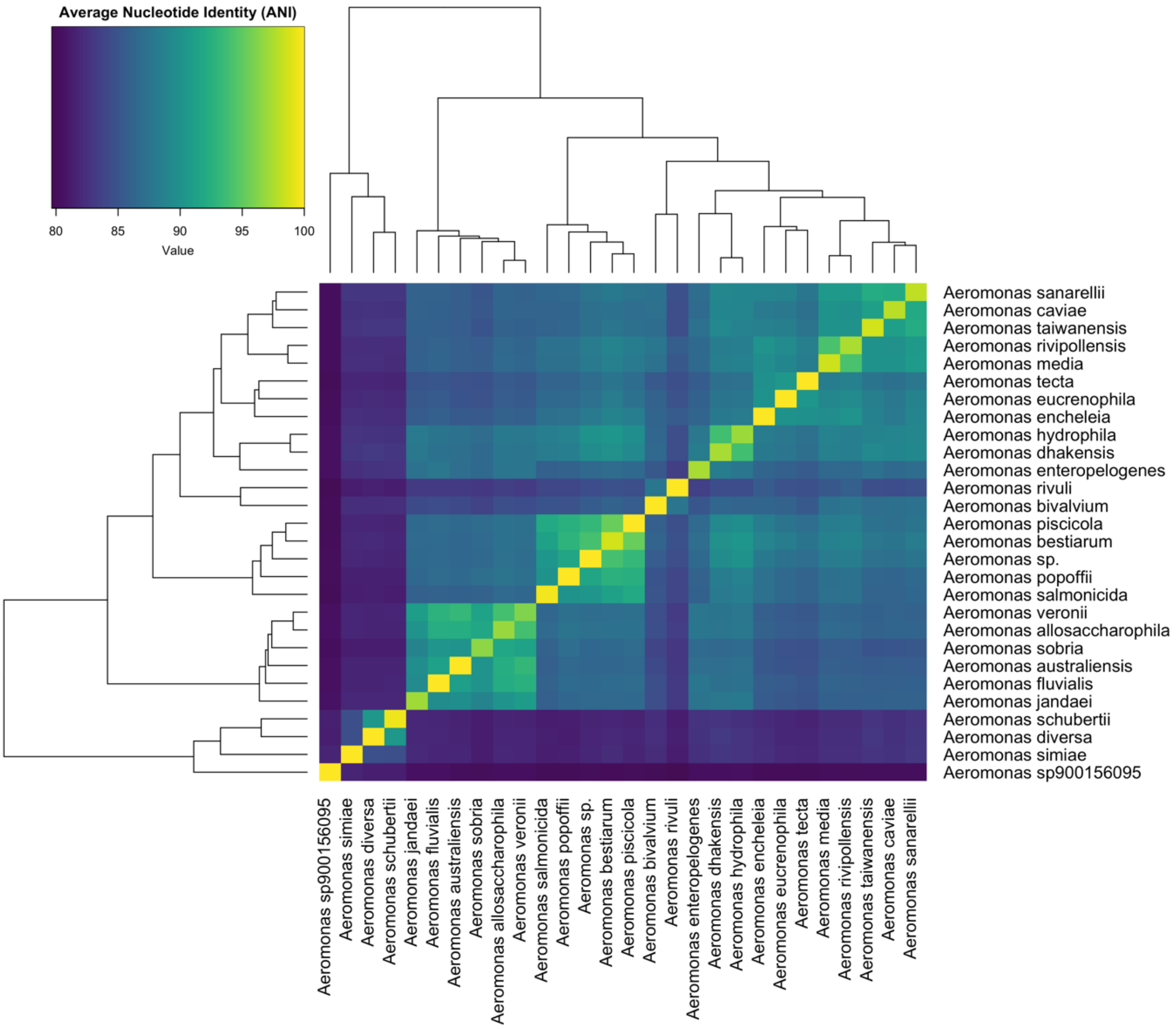


**Supplementary Figure 2: The Average Nucleotide Identity (ANI) among 1,853 *Aeromonas* spp. genomes**. The colour-coded heatmap illustrates the pairwise genomic relatedness among the genomes, based on the average ANI values. The vertical dendrogram on the sides demonstrates the hierarchical clustering of genomes according to genetic similarities. The colour scale bar indicates FastANI scores, with higher scores indicating greater similarity.

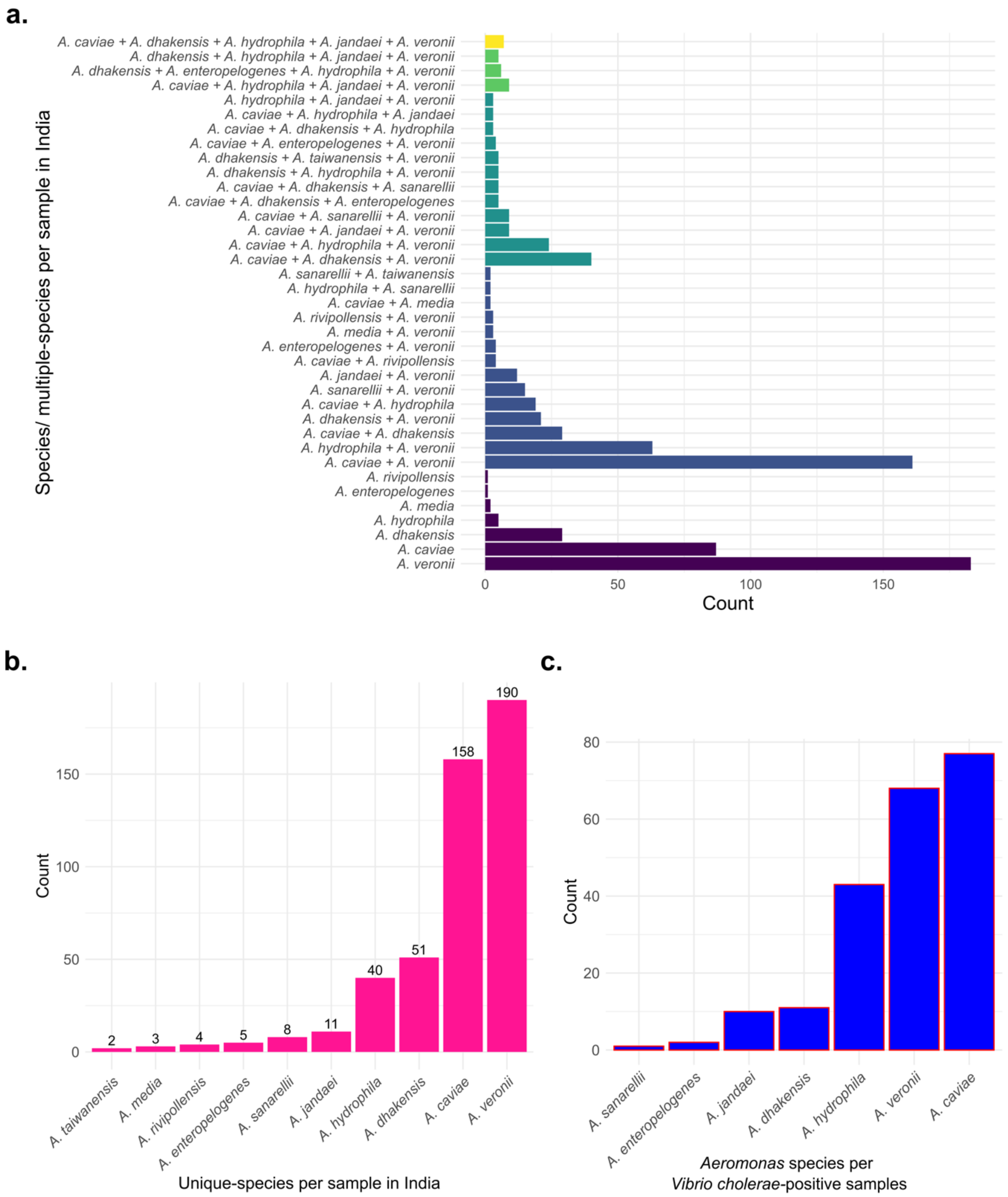


**Supplementary Figure 3: *Aeromonas* species genomes sequenced from positive water samples in Northern India. (a)** Distribution of *Aeromonas* species (y-axis) based on the presence of single or multiple species per sample (x-axis). Each bar represents a combination of species within a sample. **(b)** Abundance of *Aeromonas* species across Northern India, with one representative genome per species per sample included to account for multiple colonies sequenced per sample. Species are shown on the x-axis and their overall counts on the y-axis. **(c)** Co-occurrence of *Aeromonas* species in 91 *Vibrio cholerae*–positive samples, with species on the x-axis and their counts on the y-axis.

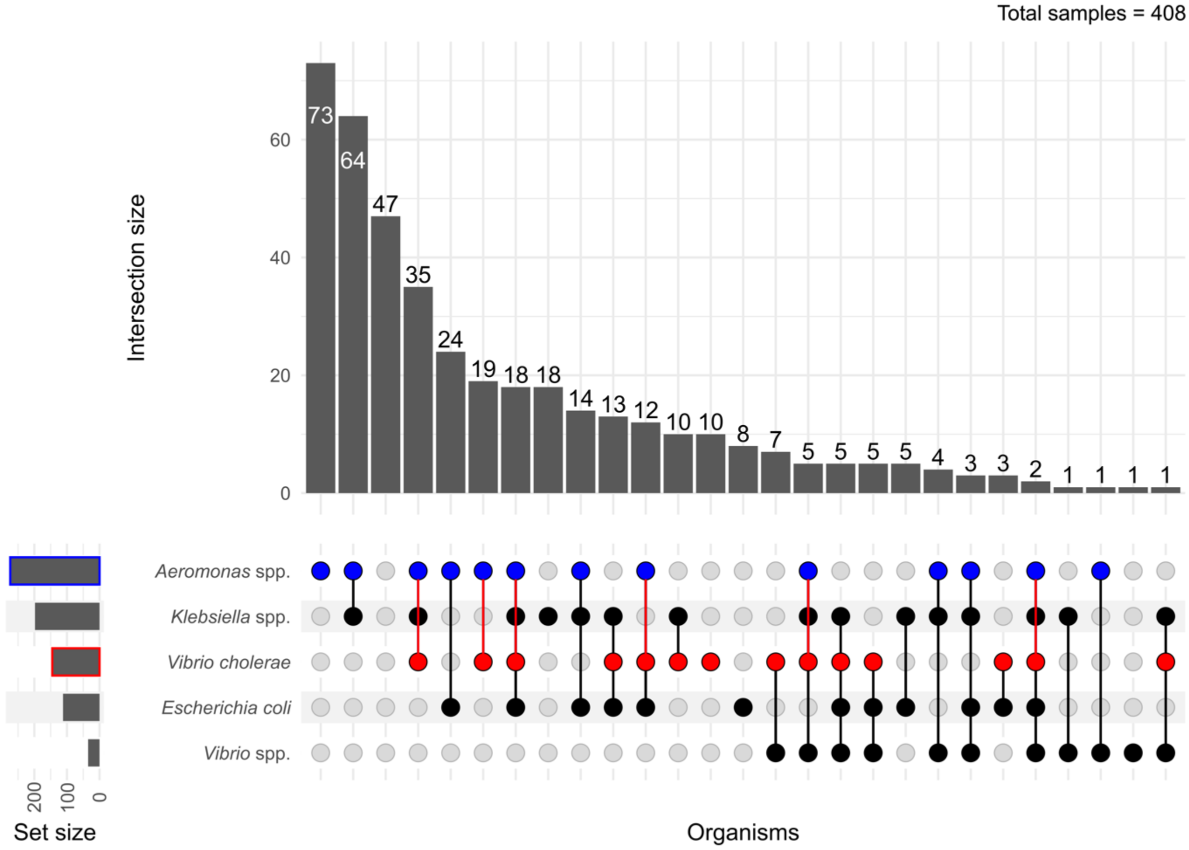


**Supplementary Figure 4:** Co-occurrence of different bacteria in 408 water samples from Northern India as determined by culturing them on selective media for *V. cholerae* and enteric pathogens (see method).

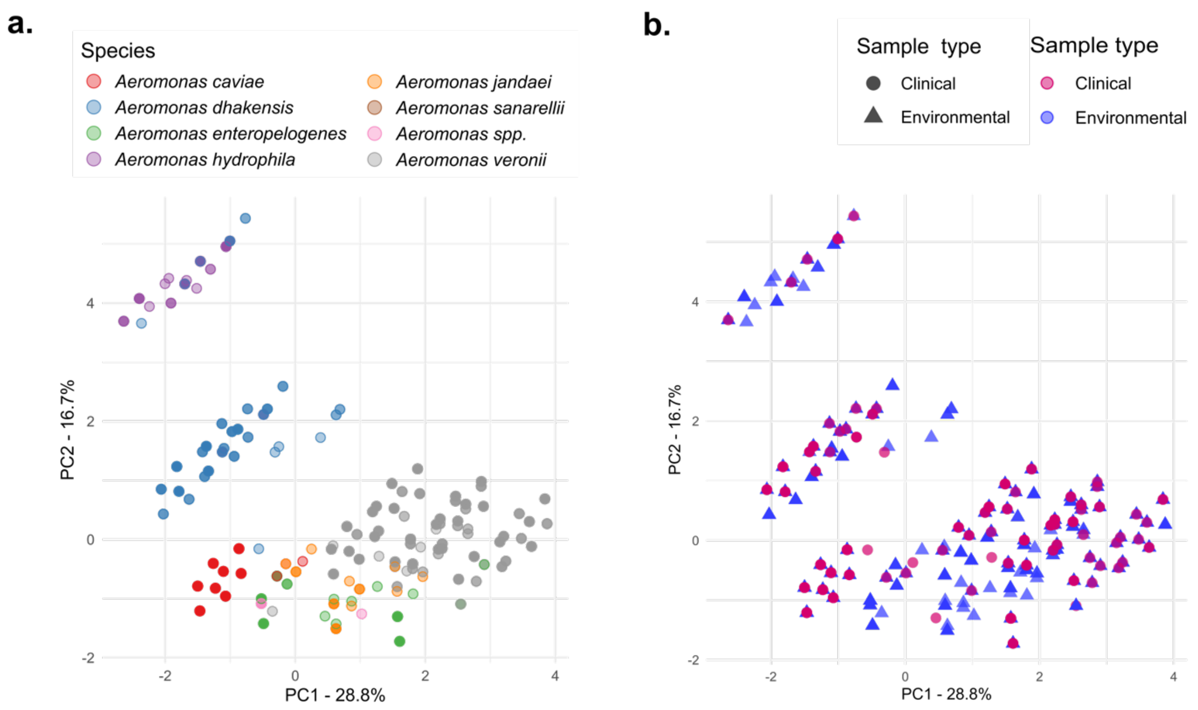


**Supplementary Figure 5: Distribution of virulence genes across 1,438 *Aeromonas* species isolates in South Asia, including Bangladesh, India, and Pakistan. (a)** Principal Component Analysis (PCA) of virulence genes across different *Aeromonas* species (see key for coloured dots representing species). **(b)** Principal Component Analysis (PCA) of virulence genes across clinical and environmental isolates (see key for coloured dots representing sample type).

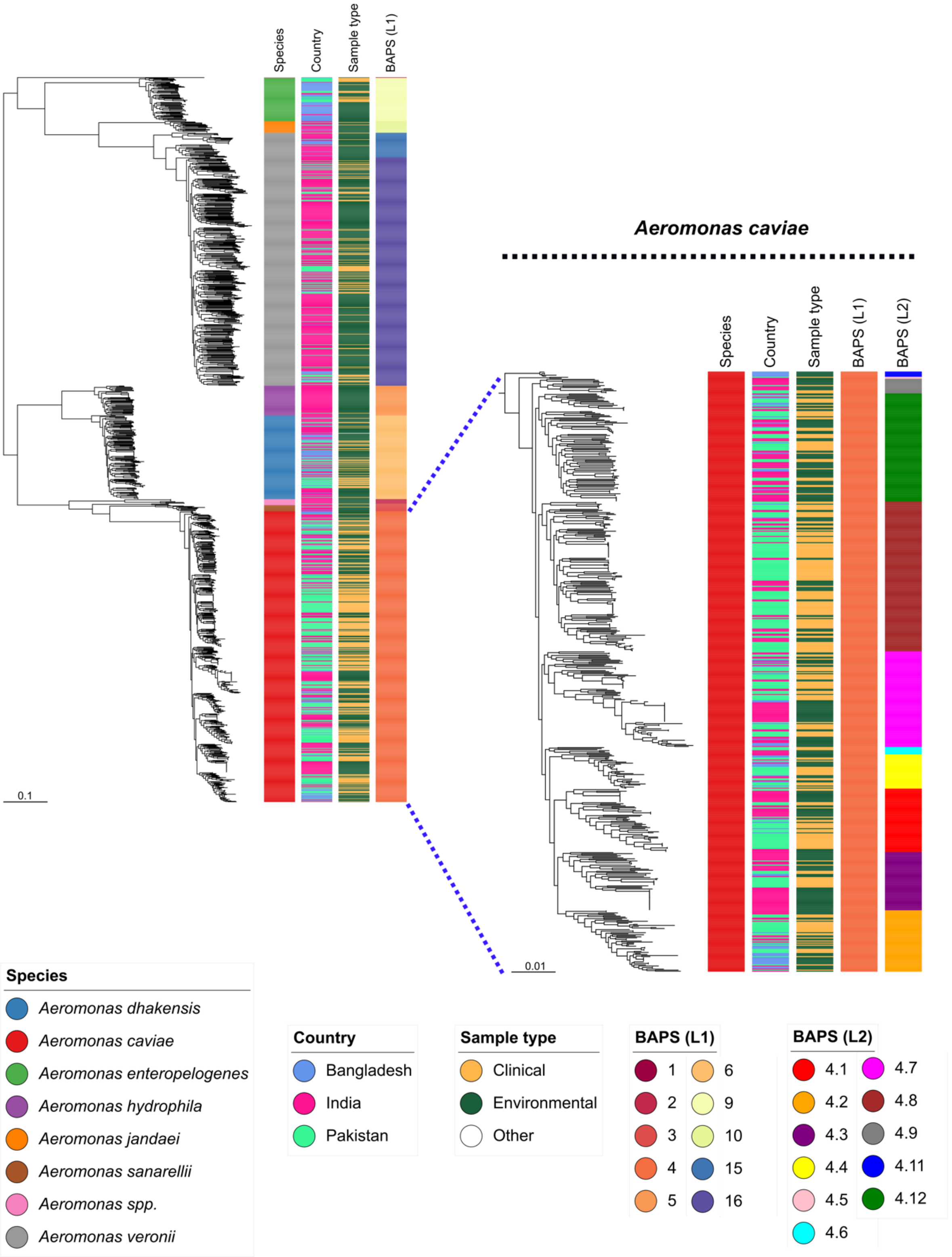


**Supplementary Figure 6: A maximum likelihood phylogenetic tree of 1,438 *Aeromonas* species genomes from South Asia (Bangladesh, India, and Pakistan), based on 2,067 core genes.** A subtree containing only *A. caviae* is highlighted separately with a dotted line. Refer to the key for interpretation of the colour strips alongside. The scale bars indicate an evolutionary distance of 0.1 nucleotide substitutions per site for the whole tree and 0.01 substitutions per site for the *A. caviae* subtree.

#### SUPPLEMENTARY DATA (TABLES)

**Supplementary Table 1:** Description of *Aeromonas* genomes by country included in this study.

| Country | Number of genomes | Study |
|---|---|---|
| Afghanistan | 2 | Blackwell, G. A. *et al.*, 2021 |
| Bangladesh | 198 | This study |
| Brazil | 1 | Blackwell, G. A. *et al.*, 2021 |
| Canada | 5 | Blackwell, G. A. *et al.*, 2021 |
| China | 62 | Blackwell, G. A. *et al.*, 2021 |
| Denmark | 100 | Blackwell, G. A. *et al.*, 2021 |
| France | 10 | Blackwell, G. A. *et al.*, 2021 |
| Germany | 1 | Blackwell, G. A. *et al.*, 2021 |
| India | 1 | Blackwell, G. A. *et al.*, 2021 |
| India | 798 | This study |
| Italy | 1 | Blackwell, G. A. *et al.*, 2021 |
| Martinique | 3 | Blackwell, G. A. *et al.*, 2021 |
| Mexico | 1 | Blackwell, G. A. *et al.*, 2021 |
| Pakistan | 441 | Klemm, E. J. *et al.*, 2024 |
| Spain | 1 | Blackwell, G. A. *et al.*, 2021 |
| Switzerland | 1 | Blackwell, G. A. *et al.*, 2021 |
| Tanzania | 2 | Blackwell, G. A. *et al.*, 2021 |
| Thailand | 1 | Blackwell, G. A. *et al.*, 2021 |
| United Kingdom | 2 | Blackwell, G. A. *et al.*, 2021 |
| United State | 71 | Blackwell, G. A. *et al.*, 2021 |
| Unknown | 151 | Blackwell, G. A. *et al.*, 2021 |

**Supplementary Table 2:** BAPS clusters and corresponding *Aeromonas* species.

| BAPS clusters | Organism |
|---|---|
| Cluster-1 | *Aeromonas schubertii, Aeromonas simiae, Aeromonas rivuli, Aeromonas diversa, Aeromonas bivalvium, Aeromonas sp900156095* |
| Cluster-2 | *Aeromonas rivipollensis, Aeromonas media, Aeromonas tecta, Aeromonas eucrenophila, Aeromonas encheleia* |
| Cluster-3 | *Aeromonas sanarellii, Aeromonas taiwanensis* |
| Cluster-4 | *Aeromonas caviae* |
| Cluster-5 | *Aeromonas hydrophila* |
| Cluster-6 | *Aeromonas dhakensis* |
| Cluster-7 | *Aeromonas popoffii, Aeromonas piscicola, Aeromonas bestiarum, Aeromonas sp.* |
| Cluster-8 | *Aeromonas salmonicida* |
| Cluster-9 | *Aeromonas enteropelogenes* |
| Cluster-10 | *Aeromonas jandaei* |
| Cluster-11 | *Aeromonas sobria* |
| Cluster-12 | *Aeromonas fluvialis* |
| Cluster-13 | *Aeromonas allosaccharophila* |
| Cluster-14 | *Aeromonas australiensis* |
| Cluster-15 | *Aeromonas veronii* |
| Cluster-16 | *Aeromonas veronii* |

**Supplementary Table 3:** *Aeromonas caviae* sublineages (BAPS) by sample type in South Asia.

| BAPS_L2 | Clinical (%) | Environmental (%) |
|---|---|---|
| Cluster-4.1 | 32 (11.19%) | 29 (9.97%) |
| Cluster-4.2 | 27 (9.44%) | 32 (11.00%) |
| Cluster-4.3 | 15 (5.24%) | 41 (14.09%) |
| Cluster-4.4 | 17 (5.94%) | 16 (5.50%) |
| Cluster-4.5 | 1 (0.35%) | 1 (0.34%) |
| Cluster-4.6 | 3 (1.05%) | 4 (1.37%) |
| Cluster-4.7 | 41 (14.34%) | 51 (17.53%) |
| Cluster-4.8 | 105 (36.71%) | 39 (13.40%) |
| Cluster-4.9 | 4 (1.40%) | 10 (3.44%) |
| Cluster-4.11 | 0 (0.00%) | 5 (1.72%) |
| Cluster-4.12 | 41 (14.34%) | 63 (21.65%) |
| Total | 286 | 291 |

**Supplementary Table 4**: Overview of Antimicrobial Resistance (AMR) genes identified in 1,853 *Aeromonas* species genomes.

| Drug class/ Organsim | *Aeromonas caviae* | *Aeromonas dhakensis* | *Aeromonas enteropelogenes* | *Aeromonas hydrophila* | *Aeromonas jandaei* | *Aeromonas salmonicida* | *Aeromonas sanarellii* | *Aeromonas spp.* | *Aeromonas veronii* |
|---|---|---|---|---|---|---|---|---|---|
| **Aminoglycoside** | *ANT(3'')-Iia, APH(6)-Id, APH(3'')-Ib, APH(3')-Ia, ANT(3'')-II-AAC(6')-Iid, aadA2, AAC(3)-Iie, AAC(6')-Ib10, AAC(6')-Ib9, AAC(6')-Ib7, aadA3, aadA15* | *AAC(6')-Ib9, ANT(3'')-Iia, aadA2* | *ANT(3'')-IIa* | *ANT(3'')-Iia, APH(6)-Id, APH(3'')-Ib, ANT(2'')-Ia, aadA2, APH(3')-Ia, AAC(3)-Iid, AAC(3)-IIIc, APH(3')-Iib, APH(3')-VI, AAC(6')-Iia, AAC(6')-Ib7* | _ | *ANT(3'')-Iia, APH(6)-Id, APH(3'')-Ib ,aadA7, APH(3')-Ia, AAC(3)-Iie, aadA2, AAC(6')-Ib9, AAC(3)-Iid, AAC(6')-Ia, ANT(2'')-Ia* | *ANT(3'')-Iia, APH(6)-Id, APH(3'')-Ib* | *ANT(3'')-Iia, AAC(6')-Ib9, APH(3'')-Ib, aadA10, APH(6)-Id, ANT(2'')-Ia, AAC(6')-Iia* | *aadA2, APH(3')-Ia, AAC(6')-Ib7, AAC(6')-Iic, AAC(6')-Iaf, ANT(3'')-Iia, ANT(2'')-Ia, AAC(3)-Iid, APH(3'')-Ib, APH(6)-Id* |
| **β-lactam (Carbapenem)** | _ | *cphA2, cphA7, imiH, cphA8* | _ | *CEPH-A3, cphA6, imiS, cphA7, imiH, cphA2* | *CEPH-A3, cphA4, cphA8* | *cphA5* | _ | *cphA8, cphA5, cphA6, cphA4, imiH, CEPH-A3* | *cphA4, CEPH-A3, cphA6, cphA8* |
| **β-lactam (Cephalosporin)** | *CTX-M-15* | *AQU-2, AQU-1, AQU-3* | _ | *CepS_beta-lactamase, AQU-2* | *AQU-2, AQU-1* | *CTX-M-3* | _ | *AQU-2* | *CepS_beta-lactamase* |
| **β-lactam (Cephamycin)** | *CMY-59* | _ | _ | _ | _ | *CMY-59* | *CMY-1, CMY-8* | _ | _ |
| **β-lactam (Penam)** | *CARB-3* | _ | _ | *CARB-3, CARB-12, FONA-5* | _ | _ | _ | _ | _ |
| **β-lactam (Cephalosporin and Cephamycin)** | *FOX-5* | _ | _ | _ | _ | *FOX-2, FOX-4, FOX-3* | _ | *FOX-2, FOX-7, FOX-8, FOX-1* | *FOX-3, FOX-2, FOX-7* |
| **β-lactam (Cephalosporin and Penam)** | *OXA-427, OXA-10, OXA-1, OXA-9* | *OXA-724, OXA-10* | *TRU-1, OXA-10* | *OXA-724, OXA-12, OXA-10* | *TRU-1, OXA-12* | *OXA-427, OXA-10, OXA-1* | *OXA-427, OXA-392* | *OXA-427, OXA-12, OXA-724, OXA-10, TRU-1, OXA-4, OXA-17* | *OXA-12, TRU-1, OXA-10, OXA-1* |
| **β-lactam (Cephalosporin and Monobactam)** | *VEB-9* | _ | _ | _ | _ | *VEB-1* | _ | _ | *VEB-1* |
| **β-lactam (Cephalosporin, Penam and Penem)** | _ | _ | _ | _ | _ | *SCO-1* | _ | _ | _ |
| **β-lactam (Carbapenem, Cephalosporin and Penam)** | *OXA-2, GES-14* | _ | _ | *SHV-134, GES-14* | _ | _ | _ | *GES-1, GES-14* | *SHV-134* |
| **β-lactam (Cephalosporin, Cephamycin and Penam)** | *MOX-6, MOX-5, MOX-8, MOX-2, MOX-7, MOX-4, MOX-3* | _ | _ | _ | _ | _ | _ | *MOX-9, MOX-6, MOX-2* | _ |
| **β-lactam (Carbapenem, Cephalosporin, Monobactam and Penam)** | *KPC-1* | _ | _ | *KPC-1* | _ | _ | _ | _ | _ |
| **β-lactam (Cephalosporin, Monobactam, Penam and Penem)** | *TEM-150* | _ | *TEM-1* | *TEM-26* | _ | *TEM-1* | _ | _ | *TEM-1, TEM-116, TEM-150* |
| **β-lactam (Carbapenem, Cephalosporin, Monobactam, Penam and Penem)** | *PER-3* | _ | _ | _ | _ | _ | _ | _ | _ |
| **Diaminopyrimidine** | *dfrA15, dfrA12, dfrA1, dfrA23* | *dfrA14, dfrA15, dfrA12* | *dfrA10, dfrA7, dfrA15* | *dfrA15, dfrA19, dfrA12, dfrA1* | _ | *dfrA1, dfrA14, dfrA12, dfrA5* | *dfrB5* | _ | *dfrA1, dfrA12, dfrA17, dfrA14* |
| **Fluoroquinolone** | *QnrS2, QnrVC4, QnrA1* | *QnrVC4* | *qacH, QnrVC4, QnrS2* | *QnrB5, QnrS2, QnrVC6* | _ | *QnrVC4, QnrVC6, qacH* | _ | *qacH* | *QnrB4, QnrS2, QepA2* |
| **Macrolide** | *mphA, mphE, mphF* | *mphA* | _ | *mphE, mphA* | _ | *mphE, mphA* | *EreA2* | *mphE* | *mphA, EreA2* |
| **Nucleoside** | _ | _ | _ | _ | _ | _ | _ | _ | _ |
| **Peptide** | *MCR-3, MCR-3.12, MCR-3.3* | _ | _ | *MCR-5* | *MCR-7.1* | *MCR-3, MCR-3.12* | _ | *MCR-3.6, MCR-3, MCR-3.8, MCR-7.1, MCR-3.12* | *MCR-3, MCR-3.12, MCR-3.6, MCR-3.3* |
| **Phenicol** | *floR, cmlB1, cmlA1, catB3, Vibrio_anguillarum_chloramphenicol_acetyltransferas1-651/651, catB8, catI, cmlA5 ,cmlA6, catB11, catII* | *cmlA5* | *cmlA5* | *cmlA5, catB8, catB3, catB11, cmlA6, floR* | _ | *floR, Ecol_catII, catB3, cmlA5* | *catB3, floR* | *cmlA5, catB3* | *catII, floR, cmx, cmlA5, catB3* |
| **Rifamycin** | *arr-3, arr-2* | _ | _ | *arr-3* | _ | *arr-3* | *arr-2* | _ | *arr-2, arr-3* |
| **Sulfonamide** | *sul2, sul1* | *sul1* | *sul1* | *sul1, sul2* | _ | *sul1, sul2* | *sul1, sul2* | *sul1* | *sul1, sul2* |
| **Tetracycline** | *tet(E), tet(A), tet(C), tet(G), tet(31)* | *tet(A), tet(E), tet(C)* | *tet(E), tet(A)* | *tet(A), tet(E), tet(C)* | *tet(A)* | *tet(A), tet (E)* | _ | *tet(A), tet (E)* | *tet(E), tet(A), tet(D)* |
| **Aminoglycoside, Fluoroquinolone** | *AAC(6')-Ib-cr* | | _ | *AAC(6')-Ib-cr* | _ | _ | _ | _ | *AAC(6')-Ib-cr* |
| **Aminoglycoside, β-lactam (Cephalosporin), Macrolide, Peptide, Rifamycin, Tetracycline** | _ | _ | *Kpne_KpnE* | _ | _ | _ | _ | _ | _ |
| **Fluoroquinolone, Macrolide and β-lactam (Penam)** | _ | _ | _ | _ | _ | _ | _ | *CRP* | _ |
| **Lincosamide, Macrolide, Oxazolidinone, Phenicol, Pleuromutilin, Streptogramin, Tetracycline** | *msrE* | _ | _ | *msrE* | _ | *msrE* | _ | *msrE* | _ |
| **Aminocoumarin, Aminoglycoside, β-lactam (Cephalosporin, Penem), Diaminopyrimidine, Fluoroquinolone, Macrolide, Phenicol and Tetracycline** | *MexD* | _ | _ | *MexD* | _ | *MexD* | _ | _ | _ |

**Supplementary Table 5**: Description of the identified β-lactamase genes in 1,853 *Aeromonas* species genomes.

| Ambler class | Genes | *A. caviae* (*n* = 608) | *A. dhakensis* (*n* = 186) | *A. enteropelogenes* (*n* = 105) | *A. hydrophila* (*n* = 124) | *A. jandaei* (*n* = 25) | *A. salmonicida* (*n* = 176) | *A. sanarellii* (*n* = 16) | *A. veronii* (*n* = 566) | *Aeromonas* spp. (*n* = 47) |
|---|---|---|---|---|---|---|---|---|---|---|
| Class A | *CARB-12* | 0 | 0 | 0 | 1 | 0 | 0 | 0 | 0 | 0 |
| | *CARB-3* | 1 | 0 | 0 | 1 | 0 | 0 | 0 | 0 | 0 |
| | *CTX-M-15* | 1 | 0 | 0 | 0 | 0 | 0 | 0 | 0 | 0 |
| | *CTX-M-3* | 0 | 0 | 0 | 0 | 0 | 1 | 0 | 0 | 4 |
| | *FONA-5* | 0 | 0 | 0 | 1 | 0 | 0 | 0 | 0 | 0 |
| | *GES-1* | 0 | 0 | 0 | 0 | 0 | 0 | 0 | 0 | 1 |
| | *GES-14* | 1 | 0 | 0 | 1 | 0 | 0 | 0 | 0 | 1 |
| | *KPC-1* | 6 | 0 | 0 | 9 | 0 | 0 | 0 | 0 | 0 |
| | *PER-3* | 4 | 0 | 0 | 0 | 0 | 0 | 0 | 5 | 0 |
| | *RSA-1* | 11 | 7 | 0 | 0 | 0 | 0 | 0 | 2 | 0 |
| | *SCO-1* | 0 | 0 | 0 | 0 | 0 | 1 | 0 | 0 | 0 |
| | *SHV-134* | 0 | 0 | 0 | 1 | 0 | 0 | 0 | 1 | 0 |
| | *TEM-1* | 0 | 0 | 1 | 0 | 0 | 1 | 0 | 4 | 4 |
| | *TEM-116* | 0 | 0 | 0 | 0 | 0 | 0 | 0 | 1 | 0 |
| | *TEM-150* | 1 | 0 | 0 | 0 | 0 | 0 | 0 | 1 | 0 |
| | *TEM-26* | 0 | 0 | 0 | 3 | 0 | 0 | 0 | 0 | 0 |
| | *VEB-1* | 5 | 0 | 0 | 0 | 0 | 1 | 0 | 13 | 0 |
| | *VEB-2* | 0 | 0 | 0 | 0 | 0 | 0 | 0 | 1 | 0 |
| | *VEB-9* | 9 | 0 | 0 | 0 | 0 | 0 | 0 | 4 | 0 |
| Class B | *CEPH-A3* | 0 | 4 | 0 | 30 | 10 | 0 | 0 | 362 | 4 |
| | *cphA2* | 0 | 72 | 0 | 31 | 0 | 0 | 0 | 0 | 0 |
| | *cphA4* | 0 | 0 | 0 | 0 | 5 | 0 | 0 | 167 | 5 |
| | *cphA5* | 0 | 0 | 0 | 0 | 0 | 113 | 0 | 0 | 2 |
| | *cphA6* | 0 | 0 | 0 | 2 | 0 | 0 | 0 | 28 | 1 |
| | *cphA7* | 0 | 41 | 1 | 25 | 0 | 0 | 0 | 0 | 0 |
| | *cphA8* | 0 | 19 | 0 | 0 | 10 | 0 | 0 | 4 | 3 |
| | *imiH* | 0 | 50 | 0 | 32 | 0 | 0 | 0 | 1 | 2 |
| | *imiS* | 0 | 0 | 0 | 2 | 0 | 0 | 0 | 0 | 0 |
| | *VIM-6* | 1 | 0 | 0 | 0 | 0 | 0 | 0 | 0 | 0 |
| Class C | *AQU-1* | 0 | 62 | 0 | 0 | 4 | 0 | 0 | 0 | 0 |
| | *AQU-2* | 0 | 49 | 0 | 24 | 16 | 0 | 0 | 1 | 0 |
| | *AQU-3* | 0 | 75 | 0 | 0 | 0 | 0 | 0 | 1 | 0 |
| | *CepS* | 0 | 0 | 0 | 100 | 0 | 0 | 0 | 14 | 0 |
| | *CMY-1* | 0 | 0 | 0 | 0 | 0 | 0 | 6 | 0 | 0 |
| | *CMY-2* | 0 | 0 | 0 | 0 | 0 | 0 | 0 | 1 | 0 |
| | *CMY-59* | 46 | 0 | 2 | 0 | 0 | 1 | 1 | 4 | 0 |
| | *CMY-8* | 0 | 0 | 0 | 0 | 0 | 0 | 10 | 0 | 0 |
| | *DHA-1* | 1 | 0 | 0 | 0 | 0 | 0 | 0 | 1 | 0 |
| | *FOX-1* | 0 | 0 | 0 | 0 | 0 | 0 | 0 | 1 | 1 |
| | *FOX-2* | 0 | 0 | 0 | 0 | 0 | 168 | 0 | 1 | 8 |
| | *FOX-3* | 0 | 0 | 0 | 0 | 0 | 3 | 0 | 21 | 0 |
| | *FOX-4* | 0 | 0 | 0 | 0 | 0 | 5 | 0 | 0 | 0 |
| | *FOX-5* | 1 | 0 | 0 | 0 | 0 | 0 | 0 | 0 | 0 |
| | *FOX-7* | 0 | 0 | 0 | 0 | 0 | 0 | 0 | 9 | 2 |
| | *FOX-8* | 0 | 0 | 0 | 0 | 0 | 0 | 0 | 0 | 1 |
| | *MOX-2* | 7 | 0 | 0 | 0 | 0 | 0 | 0 | 0 | 1 |
| | *MOX-3* | 55 | 0 | 0 | 0 | 0 | 0 | 0 | 0 | 0 |
| | *MOX-4* | 13 | 0 | 0 | 0 | 0 | 0 | 0 | 0 | 0 |
| | *MOX-5* | 60 | 0 | 0 | 0 | 0 | 0 | 0 | 0 | 0 |
| | *MOX-6* | 342 | 0 | 0 | 0 | 0 | 0 | 0 | 4 | 4 |
| | *MOX-7* | 54 | 0 | 0 | 0 | 0 | 0 | 0 | 0 | 0 |
| | *MOX-8* | 68 | 0 | 0 | 0 | 0 | 0 | 0 | 0 | 0 |
| | *MOX-9* | 0 | 0 | 0 | 0 | 0 | 0 | 0 | 0 | 15 |
| | *TRU-1* | 0 | 0 | 105 | 0 | 1 | 0 | 0 | 5 | 0 |
| Class D | *OXA-1* | 1 | 0 | 0 | 0 | 0 | 27 | 0 | 1 | 0 |
| | *OXA-10* | 12 | 2 | 8 | 1 | 0 | 25 | 0 | 11 | 1 |
| | *OXA-181* | 5 | 0 | 0 | 0 | 0 | 0 | 0 | 0 | 0 |
| | *OXA-12* | 0 | 0 | 0 | 2 | 25 | 0 | 0 | 565 | 13 |
| | *OXA-17* | 0 | 0 | 0 | 0 | 0 | 0 | 0 | 0 | 1 |
| | *OXA-2* | 14 | 0 | 0 | 0 | 0 | 0 | 0 | 0 | 0 |
| | *OXA-392* | 0 | 0 | 0 | 0 | 0 | 0 | 1 | 0 | 0 |
| | *OXA-21* | 0 | 0 | 0 | 0 | 0 | 0 | 0 | 1 | 0 |
| | *OXA-4* | 0 | 0 | 0 | 0 | 0 | 0 | 0 | 0 | 1 |
| | *OXA-427* | 608 | 0 | 0 | 0 | 0 | 176 | 16 | 0 | 26 |
| | *OXA-724* | 0 | 186 | 0 | 122 | 0 | 0 | 0 | 0 | 4 |
| | *OXA-9* | 2 | 0 | 0 | 0 | 0 | 0 | 0 | 0 | 0 |

**Supplementary Table 6:** Description of the primer sequence used for *in silico* PCR.

| **Virulence gene** | **Primer sequence (5′→3′)** | **Reference** |
|---|---|---|
| *ast* | F: ATGCACGCACGTACCGCC<br>R: ATATCCGGTCG TCG CTCTTG GT | Nhinh *et al.*, 2021 |
| *lip* | F: ATCTTCTCCGACTGGTTCGG<br>R: CCGTGCCAGGACTGGGTCTT | Kingombe *et al.*, 1999 |
| *ela* | F: ACACGGTCAAGGAGATCAAC<br>R: CGCTGGTGTTGGCCAGCAGG | Kingombe *et al.*, 1999 |
| *act* | F: GAGAAGGTGACCACCAAGAACA<br>R: AACTGACATCGGCCTTGAACTC | Kingombe *et al.*, 1999 |
| *alt* | F: GCACGGCGTGACTTCGGTGA<br>R: ACCGCGGTCTTGCAGTTGGG | Kingombe *et al.*, 1999 |
| *aerA* | F: AACCGAACTCTCCAT<br>R: CGCCTTGTCCTTGTA | Nhinh *et al.*, 2021 |
| *hlyA* | F: GGCCGGTGGCCCGAAGATACGGG<br>R: GGCGGCGCCGGACGAGACGGG | Nhinh *et al.*, 2021 |
| *fla* | F: TCCAACCGTYTGACCTC<br>R: GMYTGGTTGCGRATGGT | Nhinh *et al.*, 2021 |
| *laf* | F: GGTCTGCGCATCCAACTC<br>R: GCTCCAGACGGTTGATG | Kingombe *et al.*, 1999 |
| *ascF-G* | F: ATGAGGTCATCTGCTCGCGC<br>R: GGAGCACAACCATGGCTG AT | Kingombe *et al.*, 1999 |
